# The polyploid genome of the mitotic parthenogenetic root-knot nematode *Meloidogyne enterolobii*

**DOI:** 10.1101/586818

**Authors:** Georgios D. Koutsovoulos, Marine Poullet, Abdelnaser El Ashry, Djampa K. Kozlowski, Erika Sallet, Martine Da Rocha, Cristina Martin-Jimenez, Laetitia Perfus-Barbeoch, Juerg-Ernst Frey, Christian Ahrens, Sebastian Kiewnick, Etienne G.J. Danchin

## Abstract

Root-knot nematodes (genus *Meloidogyne*) are plant parasitic species that cause huge economic loss in the agricultural industry and affect the prosperity of communities in developing countries. Control methods against these plant pests are sparse and the current preferred method is deployment of plant cultivars bearing resistance genes against *Meloidogyne* species. However, some species such as *M. enterolobii* are not controlled by the resistance genes deployed in the most important crop plants cultivated in Europe. The recent identification of this species in Europe is thus a major concern. Like the other most damaging Meloidogyne species (e.g. *M. incognita*, *M. arenaria* and *M. javanica*), *M. enterolobii* reproduces by obligatory mitotic parthenogenesis. Genomic singularities such as a duplicated genome structure and a relatively high proportion of transposable elements have previously been described in the above mentioned mitotic parthenogenetic Meloidogyne.

To gain a better understanding of the genomic and evolutionary background we sequenced the genome of *M. enterolobii* using high coverage short and long read technologies. The information contained in the long reads helped produce a highly contiguous genome assembly of *M. enterolobii*, thus enabling us to perform high quality annotations of coding and non-coding genes, and transposable elements.

The genome assembly and annotation reveals a genome structure similar to the ones described in the other mitotic parthenogenetic Meloidogyne, described as recent hybrids. Most of the genome is present in 3 different copies that show high divergence. Because most of the genes belong to these duplicated regions only few gene losses took place, which suggest a recent polyploidization. The most likely hypothesis to reconcile high divergence between genome copies despite few gene losses and translocations is also a recent hybrid origin. Consistent with this hypothesis, we found an abundance of transposable elements at least as high as the one observed in the mitotic parthenogenetic nematodes *M. incognita* and *M. javanica*.

## Introduction

The root-knot nematode *Meloidogyne enterolobii* (syn. *M. mayaguensis*; Karssen et al. 2012) is a polyphagous species, attacking an extremely wide range of host plants including ornamentals and important agricultural crops (EPPO 2014). *M. enterolobii* is considered one of the most pathogenic and virulent root-knot nematode species, as it is able to develop and reproduce on host plants carrying resistance to the major tropical root-knot nematode species (Brito et al. 2007; Kiewnick et al. 2009; Kiewnick et al. 2011). Recent findings of *M. enterolobii* damaging cotton and soybean in the United States (Ye et al. 2013), causing detrimental effects for watermelon production areas in Mexico (Ramirez-Suarez et al. 2014), and potato in Africa (Onkendi and Moleleki 2013) demonstrate that this species spreads further. In addition, the ability of *M. enterolobii* to reproduce on plants with known sources of resistance to root-knot nematodes makes the management of this species difficult.

In 2010, *M. enterolobii* was added to the European Plant Protection Organisation (EPPO) A2 list and is now recommended for regulation as a quarantine species (Castagnone-Sereno 2012). Based on the widespread distribution and potential to establish in the Mediterranean and subtropical regions, several countries now have designated *M. enterolobii* as a quarantine pest (Elling 2013; Kiewnick et al. 2014). Currently, only a few examples for potential sources of resistance such as the *Ma* gene from the Myrobalan plum (*Prunus cerasifera*) with complete spectrum resistance against *Meloidogyne* spp. (Claverie et al., 2011) or in guava (*Psidium* spp.) and pepper accessions were reported (Freitas et al., 2014; Goncalves et al., 2014). However, these genetic resources were only tested against a single regional *M. enterolobii* population and not yet against the full range of isolates from various sources and host plants.

*M. enterolobii* is described as a species reproducing via mitotic parthenogenesis (obligatory asexual reproduction). The most damaging root-knot nematodes to worldwide agriculture (*M. incognita*, *M. javanica*, *M. arenaria*) have the same reproductive system. Given the absolute dominance of sexual reproduction in animals, parasitic success despite the absence of sex in some RKN species has been considered an evolutionary paradox (Castagnone-Sereno and Danchin 2014). The genomes of these three above-mentioned tropical RKN revealed some singularities. All the three are polyploid with highly diverged genome copies most likely resulting from hybridization events (Blanc-Mathieu et al. 2017). It was shown that this peculiar genome structure provided these species with diverged homoeologous gene copies that presented different gene expression patterns. The genomes of these three tropical RKN was also shown to be richer in transposable elements than the one of their meiotic relative *M. hapla*. These two singularities were proposed to be involved in the adaptive potential of these species despite their asexual reproduction. Because *M. enterolobii* belongs to the same Clade I (De Ley et al. 2002) as the most damaging RKN species and has the same reproductive mode, it is of great interest to explore its genome and determine similarities and differences between the species of the same clade.

In this study, we used both PacBio long reads and Illumina short read sequences to assemble the genome of *M. enterolobii*. We annotated the genome for coding and non-coding genes as well as transposable elements to determine the ploidy level, the degree of divergence between genome copies and TE abundance. Providing a solid reference genome for this species will be an important resource for further analyses with important agro-economic implications. This will pave the way for future population genomics analysis aiming at estimating the level of variability between different isolates.

## Material & Methods

### Nematode collection and DNA / RNA extraction

For genome and transcriptome sequencing, the Swiss *M. enterolobii* population was originally isolated from infected tomato root-stock obtained from an organic farm (Kiewnick et al. 2008). The population was maintained in a greenhouse at 25°C and 16h supplemental light on the cultivar Oskar F1 carrying the Mi resistance gene. The purity of the population and correct species was confirmed by species-specific PCR and DNA barcoding (Tigano et al. 2010; Kiewnick et al. 2014). For estimation of total nuclear DNA content, we used the strain Godet (from Guadeloupe France; N°75 in IPN-ISA collection).

To prepare the nematode material, heavily galled tomato roots were collected and carefully washed free from soil. Through incubation in a mist chamber, freshly-hatched second-stage juveniles (J2) were collected for two weeks and used for DNA (PacBio long reads and Illumina short read sequencing) or RNA (illumina short read) extraction. For eggs, three galled tomato roots were extracted using the NaOCl method (Hussey and Janssen 2002). In order to obtain pure egg suspensions, density-gradient centrifugation according to (Schaad and Walker 1975) was used to separate eggs from organic debris. All J2 and egg suspensions were checked using a microscope and any impurities were removed before DNA or RNA extraction.

After collecting eggs and J2s, samples were frozen in liquid nitrogen. Total DNA was extracted using the DNeasy Blood & Tissue Kit (Qiagen, The Netherlands), while total RNA was extracted using the Nucleospin RNA plant kit with the protocol modified for maximum yield (Macherey-Nagel, Oensingen, Switzerland). RNA concentration was determined with a Qubit 3.0 (Thermofisher).

### Genome and transcriptome sequencing

The Swiss *M. enterolobii* population long-insert DNA libraries were sequenced with the PacBio RS technology at the FGCZ (ETH, Switzerland) sequencing center and were complemented by additional PacBio and Illumina short read data at GATC Biotech company. DNA, extracted from nematode egg suspensions, were used to generate DNA libraries that were subsequently size-selected with the BluePippin approach, and then sequenced using the latest P6C4 chemistry, which allowed us to obtain long DNA reads of up to 35kb in length, an average length around 7 kb, and a total of 12 Gigabyte of long read sequence data. Transcriptomic data was also produced to be used as a source of evidence for gene prediction. A total of one μg RNA each was used for sequencing with the Illumina MiSeq reagent kit V3 with 2 × 300bp paired end reads.

### Experimental determination of nuclear DNA content

Flow cytometry was used to perform accurate measurement of cells DNA contents in *M. enterolobii* compared to internal standards with known genome sizes: *Caenorhabditis elegans* strain Bristol N2 (approximately 200 Mb at diploid state) and *Drosophila melanogaster* strain Cantonese S. (approximately 350 Mb at diploid state) as previously described (Blanc-Mathieu et al. 2017). Briefly, nuclei were extracted from two hundred thousand J2 infective juvenile larvae as described in (Perfus-Barbeoch et al. 2014) and stained with 75 μg/mL propidium iodide and 50 μg/mL DNAse-free RNAse. Flow cytometry analyses were carried out using an LSRII / Fortessa (BD Biosciences) flow cytometer operated with FACSDiva v6.1.3 (BD Biosciences) software. The DNA contents of the *M. enterolobii* samples were calculated by averaging the values obtained from three biological replicates.

### Genome assembly

Pacbio reads were corrected and trimmed using sprai with default parameters (http://zombie.cb.k.u-tokyo.ac.jp/sprai/). The trimmed reads were then used as input to canu assembler (Koren et al. 2017) for a first preliminary assembly. This assembly was then used to check for contamination with the blobtools pipeline (Kumar et al. 2013; Laetsch and Blaxter 2017). Briefly, Illumina libraries from both this study and (Szitenberg et al. 2017) were mapped to the genome with bwa (Li and Durbin 2009), and each contig was given a taxonomy affiliation based on BLAST results against NCBI nr database. Contigs that had coverage only in one Illumina library, GC percentage outside of the range of the estimated *M. enterolobii* GC content, and affiliation to different taxa, were considered possible contaminants. Careful investigation of each such contig was then employed to limit the false positive rate. After, only the Pacbio reads that mapped to the clean contigs were retained, and a new assembly was created with Canu. Similarly as above, this assembly was checked for contamination, and a really small percentage of contigs that was identified as contamination was removed. The clean assembly was then corrected with pilon (Walker et al. 2014) which resulted in the final frozen assembly that was used for downstream analyses.

### Completeness assessment

To assess the completeness of the genome assembly, we ran CEGMA 2.5 (Parra et al. 2007) and BUSCO v3.0 (Waterhouse et al. 2018) with the Eukaryotic dataset and *C. elegans* as a model for Augustus predictions (Stanke et al. 2008). Both pipelines search in genome assemblies for genes universally or largely conserved in Eukaryotes and produce reports on the number of genes found in complete, partial, single copy or duplicated versions in the genome under consideration. Although a nematode dataset exists in BUSCO, it only includes 8 genomes from three of the 12 described nematode clades (2, 8 and 9) (van Megen et al. 2009). Because the root-knot nematodes belong to Clade 12, we decided to instead focus on the Eukaryotic dataset which is more comprehensive (65 species, including the 8 nematodes). For comparison purpose, we also ran CEGMA and BUSCO with the same parameters on all the root-knot nematode genome assemblies that are publicly available.

### Transcriptome assembly

Adapters and low quality regions (Phred score <30) as well as regions that contained 2 or more consecutive ambiguous nucleotides were cropped using prinseq (Schmieder and Edwards 2011) resulting in 56,468,708 and 54,424,802 cleaned paired-end reads for J2 and Egg libraries, respectively. The cleaned reads were assembled using CLC Genomics Workbench 9.0 (https://www.qiagenbioinformatics.com/) with automatic bubble size and word size estimation, in simple contig mode, and minimum contig length of 200. The transcriptome assembly consisted of 110,068 for J2 library and 110,263 contigs for Egg library. The two assemblies were then merged and redundancy within and between the two assemblies was eliminated using Evigene, resulting in 74,634 non redundant coding transcripts.

### Gene prediction and annotation

Detection of gene models was done with the fully automated pipeline EuGene version 1.4 (http://eugene.toulouse.inra.fr/Downloads/egnep-Linux-x86_64.1.4.tar.gz; (Sallet et al. In Press). EuGene has been configured to integrate similarities with known proteins from i) Wormpep 221 (Lee et al. 2018) ii) *Meloidogyne incognita* proteome (Blanc-Mathieu et al. 2017) and iii) UniProtKB/Swiss-Prot database (UniProt Consortium 2018), release December 2013, proteins that were similar to those present in RepBase (Bao et al. 2015) were previously removed.

Three datasets of transcribed sequences of *Meloidogyne enterolobii* were aligned on the genome and used by EuGene as transcription evidence: i) the *de novo* assembly of the transcriptome ii) the egg stage RNAseq cleaned reads and iii) the J2 stage RNAseq cleaned reads. The alignments of the dataset 1 spanning 50% of the transcript length with at least 95% identity were retained. The alignments of the datasets 2 and 3 spanning 90% of the read length with 97% identity were retained.

The EuGene default configuration was edited to i) set the “preserve” parameter to 1 for all datasets ii) set “gmap_intron_filter” parameter to 1 iii) set the minimum length of introns to 35 iv) allow the non canonical donor spliced site “GC”. Finally, the Nematode specific Weight Array Method matrices were used to score the splice sites (http://eugene.toulouse.inra.fr/Downloads/WAM_nematodes_20171017.tar.gz).

### Prediction and annotation of transposable elements

Prediction and annotation of repeats were performed with the REPET meta-pipeline which regroups TEdenovo and TEannot pipelines (Flutre et al. 2011).

#### Data pre-processing

In order to improve homology search during the de-novo analysis (Frith 2011), low complexity regions and Satellites and Simple Repeats (SSR) were masked with Ns in the *M. enterolobii* genome using RepeatMasker-4.0 (-noint) (http://www.repeatmasker.org). Then, the genome was split into chunks using DBchunks (REPET tool) to remove stretches of 11 ‘N’s or more. Because small sequences can interfere with TE discovery, chunk’s L50 was calculated and used as a minimum size threshold.

#### TE de-novo prediction

Chunks with a length > L50 were used to build a TE consensus library using the TEdenovo pipeline (see TEdenovo.cfg file in supplementary for parameters values). The genome was aligned to itself using Blaster (Quesneville et al. 2003) and High Scoring segment Pairs (HSPs) were detected. The repetitive HSPs were clustered by Recon (Bao and Eddy 2002) and Grouper (Quesneville et al. 2003). A consensus sequence was created for each cluster based on a multiple alignment with MAP (Huang 1994). Consensus sequences were analysed in order to find homology with known TE and to detect TE-related features such as specific HMM-profiles and structural characteristics. Therefore, curated libraries provided by the REPET development team (URGI) were used: repbase 20.05 (aa and nt), a concatenation of pfam 27.0 and GyDB 2.0 hmm-profiles databanks and rRNA Eukaryota databank. Detected features were used by PASTEClassifier (Hoede et al. 2014) to classify consensus sequences in the different TE orders defined in Wicker’s classification (Wicker et al. 2007). SSR and under-represented unclassified consensuses were filtered out.

#### Semi-automated TE consensus filtering

The TE consensus library was filtered as follows in order to minimise false positive sequences and redundancy.A draft annotation of the whole genome based on the consensus library was performed using TEannot (steps 1, 2, 3, 7; see TEannot1.cfg file in supplementary for parameters values). Only consensuses with at least one Full-Length Copy (FLC) annotated on the genome were retained to constitute a new ‘filtered TE consensus library’. Consensuses with at least one FLC annotated over the genome were identified with PostAnalyzeTELib.py and their sequences were extracted using GetSequencesFromAnnotations.py (REPET tools).

#### TE whole genome annotation

The TEannot pipeline was used to annotate the whole *M. enterolobii* genome from the ‘filtered TE consensus library’ (steps 1, 2, 3, 4, 5, 7, 8; see TEannot2.cfg file in supplementary for parameters values). TE consensus sequences from the 'filtered TE consensus library' were aligned on the genome using Blaster, Censor (Jurka et al. 1996), and RepeatMasker. The results of the three methods were concatenated and MATCHER (Quesneville et al. 2003) was used to remove overlapping HSPs and make connections with the “join” procedure. SSRs were detected by TRF (Benson 1999), Mreps (Kolpakov et al. 2003), and RepeatMasker, and then merged. Eventually, after some redundancy removal and application of the “long join” procedure (distant fragments belonging to the same copies are joined), annotations were exported for post-processing.

#### Annotations filtering and post-processing

Using in-house Python scripts, we retrieved TE annotation with > 85 % identity to consensus sequence and length > 150 nucleotides. Moreover, only TE classified as retrotransposon (Wicker’s class I) and DNA-transposon (Wicker’s class II) were retained. This script also allowed to evaluate annotation redundancy and to describe TE coverage on the genome regarding Wicker’s classification.

### Genome structure and duplications

We used MCScanX (Wang et al. 2012) to detect and classify duplications and study the genome structure. In a first step, all the predicted protein sequences from Eugene annotation were self-blasted to determine homologous relationships between them, with an e-value threshold of 1e-10. Using homology information from the all-against-all blast output and gene location information from the GFF3 annotation file, MCScanX detects duplicated protein-coding genes and classify them in the following categories, (i) singleton when no duplicates are found in the assembly, (ii) proximal when duplicates are on the same contig and separated by 1 to 10 genes, (iii) tandem when duplicates are consecutive, (iv) whole genome duplication (WGD) or segmental when duplicates form collinear blocks with other pairs of duplicated genes, and (v) dispersed when the duplicates cannot be assigned to any of the above-mentioned categories. For detection of WGD / segmental duplications we required at least 3 collinear gene pairs.

To measure average nucleotide divergence between duplicated genomic regions (WGD/segmental), collinear blocks were aligned with nucmer (Kurtz et al. 2004). Average nucleotide divergence at the coding level was measured by aligning gene pairs present in duplicated blocks with T-Coffee (Notredame et al. 2000).

## Results

### The M. enterolobii genome is the most complete annotated one available to date

We assembled the *M. enterolobii* Swiss strain in 4,437 contigs for a total genome size of 240 Mb and a N50 length of 143 kb. This assembled genome size is in the range of the calculated nuclear DNA content *via* flow cytometry experiments (274.69±18.52 Mb, Fig 1). This suggests that most of the *M. enterolobii* genome has been captured in this assembly. The difference between the genome assembly size and the total estimated DNA content ranges from 16 to up to 53 Mb and could be due either to duplicated and repetitive genome regions that have not been correctly separated during assembly or to differences in genome sizes between the Swiss *M. enterolob*ii strain we have sequences and the ‘Guadeloupe’ strain used for DNA content measurement via flow cytometry. Overall, the genome assembly we produced constitutes a significant improvement in terms of contiguity and completeness compared the only *M. enterolobii* draft genome that was available to date (Szitenberg et al. 2017). The genome size grew from 162.4 to 240 Mb and now fits the estimated DNA content. The number of contigs / supercontigs was divided by >10 (46,090 to 4,437) while the N50 length was multiplied by >15 (9.3 to 143 kb), Table 1.

**Figure 1.**
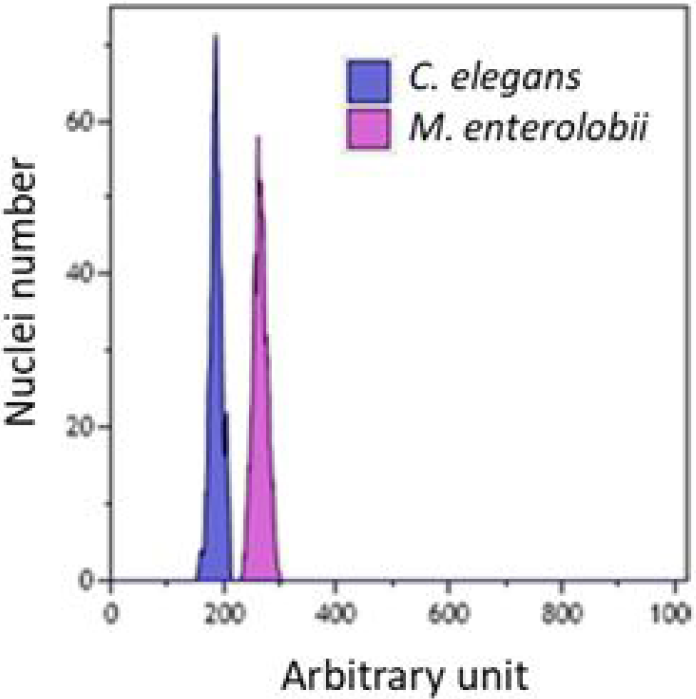
Relative DNA staining in nuclei of *M. enterolobii*. Cytogram example obtained after gating on G0/G1 nuclei (arbitrary units) from M. enterolobii when processed mixed together with C. elegans, the internal standard (diploid genome size is 200 Mb).

**Table 1.** *Meloidogyne* genome features

| Species / Feature | Assembly size (Mb) | Nuclear DNA content (Mb) | % complete CEG (C) (copies)<br>%Partial (P) | % complete BUSCO (C)<br>% fragment (F)<br>% missing (M) | N50 (kb) | # contigs/scaffolds | GC % |
| --- | --- | --- | --- | --- | --- | --- | --- |
| *M. enterolobii* (Swiss)<sup>1</sup> | 240 | 275 ±19 | C : 94.76 (3.30)<br>P : 96.77 | C:87.5%<br>F:3.6%; M:8.9% | 143 | 4,437 | 30 |
| *M. enterolobii* (L30)<sup>2</sup> | 162.4 | NA | C : 81.45 (2.66)<br>P : 89.92 | C:79.9%<br>F:8.9%; M:11.2% | 9.3 | 46,090 | 30.2 |
| *M. incognita* (Morelos, 2017)<sup>3</sup> | 183.5 | 189 ±15 | C : 94.76 (2.93)<br>P :96.77 | C:88.5%<br>F:3.0%; M:8.5% | 38.6 | 12,091 | 29.8 |
| *M. incognita* (W1)<sup>2</sup> | 122 | NA | C : 82.66 (2.34)<br>P :89.52 | C:80.2%<br>F:7.9%; M:11.9% | 16.5 | 33,735 | 30.6 |
| *M. incognita* (Morelos, 2008)<sup>4</sup> | 86 | 189 ±15 | C : 74.6 (1.77)<br>P :78.63 | C:71.3%<br>F:7.3%; M:21.4% | 82.8 | 2,817 | 31.4 |
| *M. javanica* (Avignon)<sup>3</sup> | 235.8 | 297 ±27 | C : 92.74 (3.68)<br>P :95.56 | C:90.1%<br>F:2.3%; M:7.6% | 10.4 | 31,341 | 30 |
| *M. javanica* (VW4)<sup>2</sup> | 142.6 | NA | C :89.52 (2.71)<br>P : 95.16 | C:87.5%<br>F:4.3%; M:8.2% | 14.1 | 34,394 | 30.2 |
| *M. arenaria* (Guadeloupe)<sup>3</sup> | 258.1 | 304 ±9 | C :94.76 (3.66)<br>P :95.56 | C:87.1%<br>F:4.3%; M:8.6% | 16.5 | 26,196 | 30 |
| *M. arenaria* (A2-O)<sup>5</sup> | 284.05 | NA | C : 94.76 (3.57)<br>P :96.77 | C:87.1%<br>F:2.6%; M:10.3% | 204.6 | 2,224 | 30 |
| *M. arenaria* (HarA)<sup>2</sup> | 163.8 | NA | C :91.53 (2.74)<br>P :95.97 | C : 78.2<br>F :12.2; M:9.6 | 10.5 | 46,509 | 30.3 |
| *M. hapla* (VW9)<sup>6</sup> | 53.6 | 121 ±3 | C :93.55 (1.19)<br>P : 95.56 | C:87.4%<br>F:4.3%; M:8.3% | 83.6 | 1,523 | 27.4 |
| *M. floridensis* (JB5)<sup>7</sup> | 99.9 | NA | C : 56.45 (1.95)<br>P : 76.61 | C:54.1%<br>F:28.7%; M:17.2% | 3.5 | 81,111 | 29.7 |
| *M. floridensis* (SJF1)<sup>2</sup> | 74.9 | NA | C : 77.42 (1.71)<br>P : 83.87 | C:76.5%<br>F:7.6%; M:15.9% | 13.3 | 9,134 | 30.2 |
| *M. graminicola* (IARI)<sup>8</sup> | 38.18 | NA | C : 84.27 (1.34)<br>P : 90.73 | C:73.6%<br>F:15.2%; M:11.2% | 20.4 | 4,304 | 23.05 |

<sup>1</sup>This work, <sup>2</sup>(Szitenberg et al. 2017), <sup>3</sup>(Blanc-Mathieu et al. 2017), <sup>4</sup>(Abad et al. 2008), <sup>5</sup>(Sato et al. 2018), <sup>6</sup>(Opperman et al. 2008), <sup>7</sup>(Lunt et al. 2014), <sup>8</sup>(Somvanshi et al. 2018)

To further assess genome completeness, we used CEGMA and BUSCO and this allowed to retrieve 94.76 and 87.5 % of the CEGMA and eukaryotic BUSCO genes in complete length, respectively (Table 1). The CEGMA score is the highest obtained for a *Meloidogyne* to date and is only paralleled by the 2017 versions of *M. incognita* and *M. arenaria* (Blanc-Mathieu et al. 2017) as well as the 2018 version of *M. arenaria* (Sato et al. 2018). Similarly, the BUSCO scores obtained are only surpassed by those of *M. incognita* and *M. javanica* produced in 2017 (Blanc-Mathieu et al. 2017). For comparison, the model nematode *C. elegans* reaches CEGMA and eukaryotic BUSCO completeness scores of 96.37 and 94.7, respectively. However, it should be noted that the whole protein set of *C. elegans* is part of the training set for both CEGMA and BUSCO.

Interestingly, the average number of copy per CEGMA gene in *M. enterolobii* was 3.3. This suggests that genes usually present in single copy across eukaryotes would exist in ~3 copies in *M. enterolobii*. Interestingly, the CEGMA genes are found in 2.93, 3.68 and 3.66 copies in the 2017 versions of the *M. incognita*, *M. javanica* and *M. arenaria* genomes, respectively described as triploid and degenerate tetraploids. The CEGMA copy number in *M. enterolobii* would suggest a triploid genome structure. In contrast, the genomes of *M. hapla* and *C. elegans* have an average number of CEGMA gene copies of 1.19 and 1.09, respectively which is consistent with their homozygous diploid nature and the corresponding haploid assembly.

### The majority of *M. enterolobii* genes are duplicated

Using the EUGENE pipeline, we predicted 63,841 genes, of which 59,773 were coding for proteins. The coding part spans 61.9Mb (~26%) of the genome assembly length. The GC percent was higher in the coding portion (34.3%) than in the whole genome (30.0%). Spliceosomal introns were detected in 88% of protein-coding genes, with an average of 6.2 exons per gene. Similarly to the other tropical RKN genomes, 1% of splice sites have a non-canonical GG donor dinucleotide (canonical is GT).

We aligned the CDS corresponding to the whole set of predicted proteins to the respective reference genomes of *M. enterolobii*, *M. incognita* Morelos and *M. hapla* VW9 using the procedure explained in (Blanc-Mathieu et al. 2017). In the diploid meiotic species *M. hapla*, 90% of the CDS mapped to a unique locus in the reference genome and can be considered single-copy genes. In contrast, only 12 and 13% of *M. enterolobii* and *M. incognita* CDS map a unique locus on their respective genome assembly. Hence, the vast majority of protein-coding genes are in multiple copies in *M. enterolobii* and *M. incognita* (Fig 2). Furthermore, both *M. enterolobii* and *M. incognita* show a peak of alignment at 3 different loci. This suggests a possible triploid genome structure.

**Figure 2.**
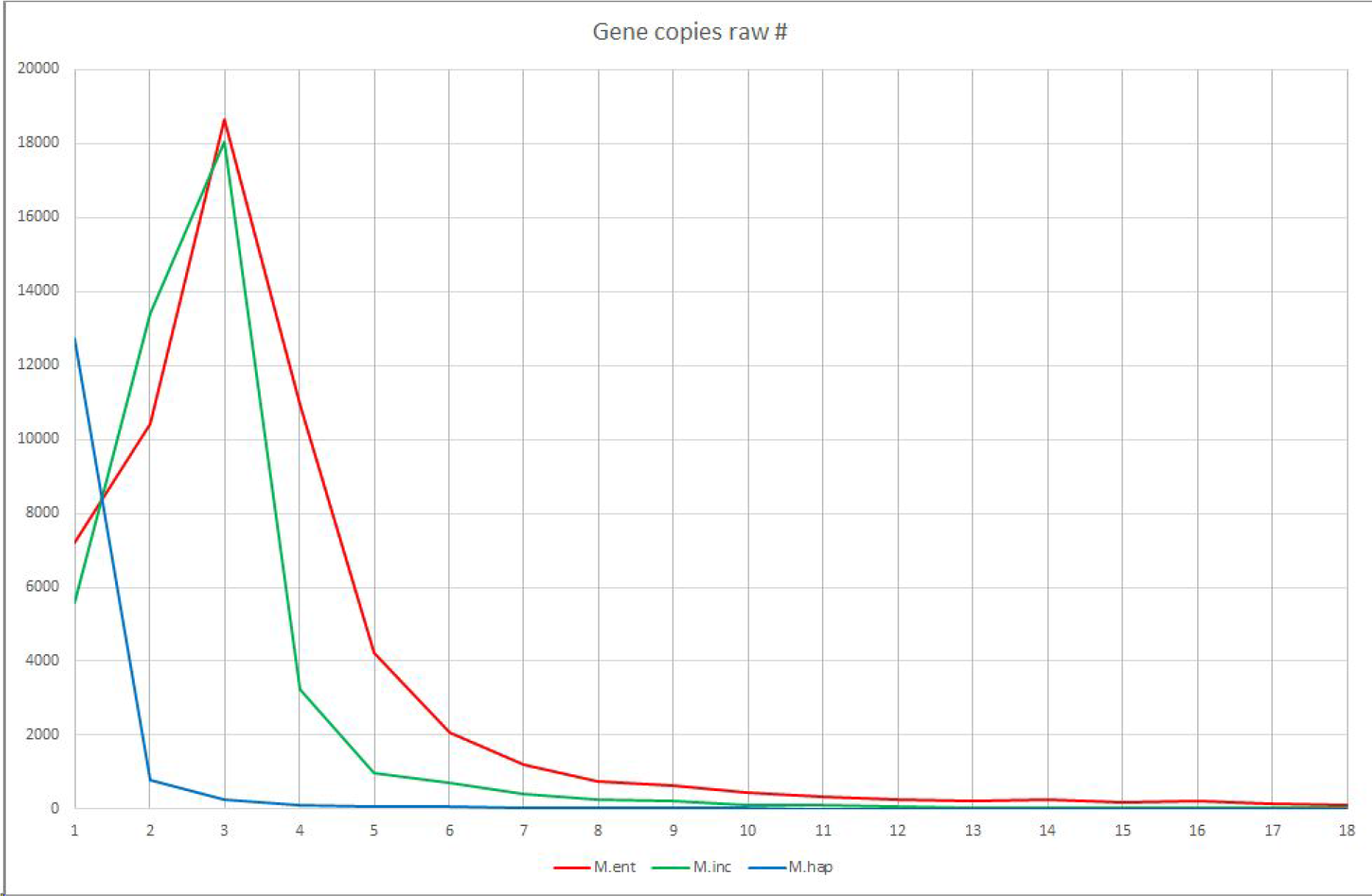
Number of CDS mapping n loci in Meloidogyne genomes. Number of CDS mapping with >95% identity on >66% on their length at n loci on the respective reference genomes.

EUGENE also predicted 4,068 non protein-coding genes (e.g. ribosomal, tRNA, splice leader genes). None of them had predicted intron but similarly to protein-coding genes the average GC content was higher (34.1%) than for the rest of the genome.

### The genome is most likely triploid

MCScanX analyses showed that around 94% of the protein-coding genes are duplicated in *M. enterolobii*. Depending on the block size parameter used in the analyses (3 to 5), the classification of the duplication differs slightly, but overall 60% of the genes are part of the whole genome duplication category, 29% of the genes have been classified as dispersed duplicates, and 5% of the genes are proximal or tandem duplicates. Selecting a cutoff of at least 3 collinear genes, we found 62% of genes are collinear forming 2,892 collinear blocks. Then, we investigated the duplication depth of genes in the collinear blocks, and similarly to previous analyses, we observed nearly half of the collinear genes to be in three copies, while the others are in various duplication depths.

Sequence divergence between gene copies showed high degrees of identity both at protein and nucleotide levels (~96%). However, the peak of the protein identity distribution is at 99% while the peak of nucleotide identity distribution is at 97%, indicating that at protein level there is higher conservation. Aligning the whole homologous duplicated blocks shows a drop in identity as expected to 92%, with the peak of distribution being at 90% (Fig 3). This is similar to the ~8% divergence measured between duplicated genome blocks in the other mitotic parthenogenetic Meloidogyne (Blanc-Mathieu et al. 2017).

**Figure 3.**
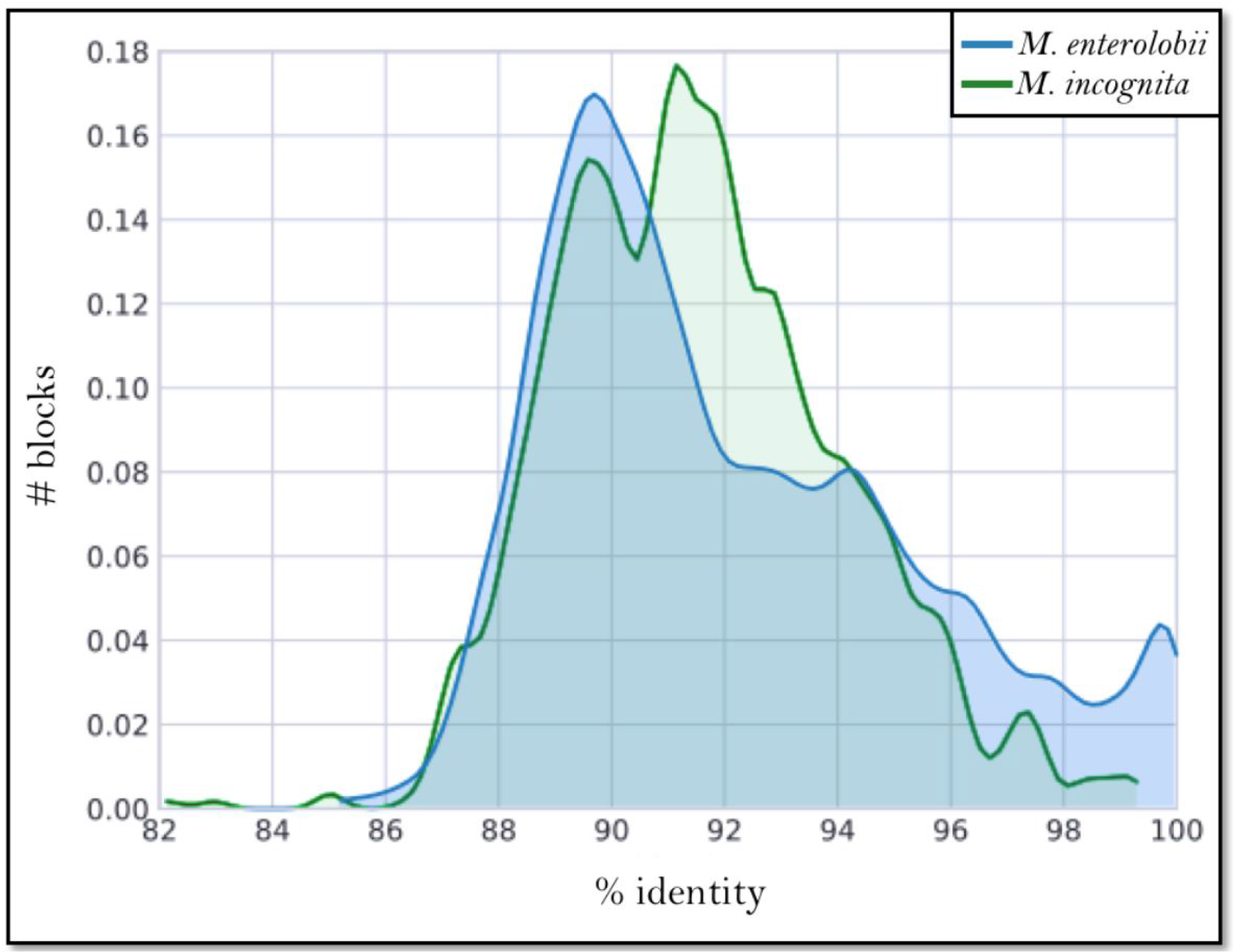
Nucleotide identity computed from NUCmer alignments within homologous duplicated blocks. The colour blue is for *M. enterolobii*, the green one for *M. incognita*. They are two apomictic nematodes species. *M. enterolobii* seems to present one bigger peak at 89.9% and two smaller at 94.2% and 99.9% of identity, whereas *M. incognita* shows almost two overlapping peaks at 89.5% and 91.4% of identity.

### The *M. enterolobii* genome is rich in non-autonomous transposable elements

Repetitive elements span 17.49% of the *M. enterolobii* genome assembly size (Table 2). Using the same protocol, we also annotated *M. incognita* and *M. javanica* genomes and repetitive elements represent 9.44% and 10.16% of their genome sizes, respectively. In *M. enterolobii,* 69.38% of the repetitive elements (12.13% of the genome size) could be assigned to Wicker's class I (4.46%) or class II (7.67%). The majority of classified TEs have been assigned to specific Orders leading to more accurate annotation.

**Table 2.** Repetitive elements content in the genome as a percentage of the genome size. Unclassified are repetitive elements that do not possess recognizable feature of either Class I (retrotransposons) or Class II (DNA transposons). CLASS_1_LIKE and CLASS_2_LIKE are repetitive elements that do possess the characteristic features of either Class I or Class II elements but could not be further classified.

|  |  | *M. incognita* | *M. javanica* | *M. enterolobii* |
| --- | --- | --- | --- | --- |
|  | assembly size (bp) | 183531997 | 235798407 | 240054310 |
|  | TE order as genome percentage |  |  |  |
| **Class I** | LTR | 1.047095 | 1.466235 | 1.825122 |
|  | DIRS | 0.038791 | 0.01735 | 0.288098 |
|  | PLE | 0 | 0 | 0 |
|  | LINE | 0.380482 | 0.266408 | 0.872724 |
|  | SINE | 0.008119 | 0.004899 | 0.002169 |
|  | TRIM | 0.206717 | 0.104754 | 0.786364 |
|  | LARD | 0.134367 | 0.111066 | 0.421383 |
|  | CLASS_1_LIKE | 0.064654 | 0.120184 | 0.264441 |
| **Class II** | TIR | 2.039291 | 1.940418 | 2.199123 |
|  | HELITRON | 0.153739 | 0.218203 | 0.613834 |
|  | MAVERICK | 1.159525 | 1.039514 | 2.916335 |
|  | MITE | 1.932049 | 1.41853 | 1.914862 |
|  | CLASS_2_LIKE | 0.010192 | 0.014453 | 0.031035 |
|  | potHostGenesOrOther | 1.641508 | 2.591996 | 4.120049 |
|  | unclassified | 0.624961 | 0.844605 | 1.234587 |
|  | total | 7.175021 | 6.722014 | 12.13549 |

Interestingly ~1/4 of the TE annotations (3.12% of the genome) correspond to non-autonomous TRIM, LARD, and MITEs TEs meaning a substantial part of the TE content in *M. enterolobii* could have been derived from decayed autonomous elements that have lost their ability to transpose by themselves in the genome. This abundance of non-autonomous TEs is due to a higher proportion of the genome occupied by TRIM and LARD elements in *M. enterolobii* than in the two other root-knot nematode genomes analyzed here (7.9 and 4.2%, respectively vs. 1.0 - 2.0% and 1.1 −1.3%, respectively for TRIM and LARD). The proportion of MITEs, however, is comparable.

## Discussion

The genome assembly and annotation of *M. enterolobii* is a first step in understanding the complex evolutionary history of this species. It will allow to test new emerging hypotheses in an ever changing environment. Furthermore, it will be a valuable resource moving forward in battling this emerging plant parasitic nematode. The high contiguity of the genome enabled us to produce a thorough gene and TE annotation which can be used alongside other root-knot nematode sequencing projects to explore further the intricacies of the genus *Meloidogyne*.

## Author contribution

EGJD, SK, GDK, DKK, MP and LPB wrote the manuscript. LPB and CMJ performed measurement of the nuclear DNA content and analysed the results. GDK assembled and cleaned the genome and participated in the analysis of the genome structure. EGJD participated in the analysis of the genome structure and performed the comparative assessment of genome. MP performed the genome structure and divergence analysis.

ES performed the gene prediction and analysed the results. DKK performed the transposable elements annotation and analysis. AEA produced the material for genome and transcriptome sequencing and analysed the results. AEA and MDR assembled the transcriptome and analysed the results. JEF, CA and SK identified, discovered and originally reared the *M. enterolobii* Swiss strain used for genome and transcriptome sequencing.

## Acknowledgements

We would like to thank Jérôme Gouzy for help and advices in gene prediction step. Julie Cazareth from “Imaging and Cytometry platform” at IPMC for tech support. GK has received the support of the EU in the framework of the Marie-Curie FP7 COFUND People Programme, through the award of an AgreenSkills+ fellowship (under grant number 609398). CMJ received funding from the ASEXEVOL project, grant number 392 ANR-13-JSV7-0006 which also funded bioinformatics equipment used for this paper.

